# A genetic perspective on the relationship between eudaimonic –and hedonic well-being

**DOI:** 10.1101/283036

**Authors:** B.M.L. Baselmans, M. Bartels

## Abstract

Whether hedonism or eudaimonism are two distinguishable forms of well-being is a topic of ongoing debate. To shed light on the relation between the two, large-scale available molecular genetic data were leveraged to gain more insight into the genetic architecture of the overlap between hedonic and eudaimonic well-being. Hence, we conducted the first genome-wide association studies (GWAS) of eudaimonic well-being (*N* = ∼108K) and linked it to a GWAS of hedonic well-being (*N = ∼* 222K). We identified the first two genome-wide significant independent loci for eudaimonic well-being and 6 independent loci for hedonic well-being. Joint analyses revealed a moderate phenotypic correlation (*r* = 0.53), but a high genetic correlation (*r_g_* = 0.78) between eudaimonic and hedonic well-being. For both traits we identified enrichment in the frontal cortex -and cingulate cortex as well as the cerebellum to be top ranked. Bi-directional Mendelian Randomization analyses using two-sample MR indicated some evidence for a causal relationship from hedonic well-being to eudaimonic well-being whereas no evidence was found for the reverse. Additionally, genetic correlations patterns with a range of positive and negative related phenotypes were largely similar for hedonic –and eudaimonic well-being. Our results reveal a large genetic overlap between hedonism and eudaimonism.

## Introduction

For centuries, people have asked themselves questions about well-being with hedonic well-being and eudaimonic well-being as its major philosophical schools of thoughts. Hedonic well-being concerns the balance of pleasure over pain, with Aristippus (c. 435 –c. 356 BCE), as one of its founders ^1^. Whereas the hedonic tradition focused on what is good for a person, the eudaimonic tradition took well-being to centre around virtuous activity, defined as knowledge (practiced over time) and the fulfilment of human capacities ^2^. One of the important founders of eudaimonic well-being is Aristotele (c. 384 – c. 322 BCE), who was a true opponent of the hedonistic school of thought describing it as “*vulgar*” ^3^. According to Aristotle, eudaimonic well-being is more than being happy and is it about the actualization of the human potential ^4^.

In contemporary behavioural and social sciences, the term hedonic well-being is used less frequently. A reason for this is that hedonism as a theoretical (data-free) concept is difficult to quantify. To redefine the hedonic line of thought in an operational construct, the subjective well-being (SWB) definition, as proposed by Diener ^5^, is widely adopted. Herein, SWB consists of three hallmarks: 1) it is subjective; 2) it includes positive measures (not just the absence of negative measures), and 3) it includes a global assessment of all aspects of a person’s life. SWB has been repeatedly found to be associated with health and mortality e.g. ^6–9^. Analogous to hedonism, the term eudaimonic well-being has gradually shifted towards psychological well-being (PWB) in contemporary science. To assess PWB, six core dimensions are widely used: self-acceptance, positive relations with others, autonomy, environmental mastery, purpose in life, and personal growth ^10^. Several studies have found that people who believe their lives have meaning or purpose appear better off, with better mental and physical health and engagement in healthier life styles ^11–16^.

Although, it is recognized that modern-day hedonism and eudaimonism are central concepts of well-being, the overlap and distinction between these two forms of well-being is a topic of an ongoing debate ^1, 17–23^. Factor analytic studies show that hedonic and eudaimonic aspects of well-being load on separate yet highly correlated factors, with correlations in the range of 0.81 to 0.92 ^24–26^. Application of less restrictive exploratory structural equation modelling, results in a correlation of 0.60 between hedonic and eudaimonic well-being ^22^. A more in-depth overview of the reported correlation between hedonic and eudaimonic uncovers a wide spread in correlations resulting from differences in degree of centrality (if the hedonic measures are the core aspect of the analyses or if the correlation is based on correlates of the concepts), application of different categories of analyses (if hedonia and eudaimonia is considered an orientation, behavior, experience, or function) and level of measurement (state versus trait) ^20^.

A way to provide more clarity on the overlap and distinction of hedonic and eudaimonic well-being is by exploring the underlying sources of overlap. Differences in both hedonic and eudaimonic well-being have been found to be partly genetic. Twin-family studies, which contrast the resemblance of monozygotic (MZ), dizygotic (DZ) twins and their non-twin siblings or other family members, report heritability estimates in the range of 30 – 64% for both hedonic and eudaimonic well-being ^27, 28^. Most molecular genetic work, so far, focused on hedonic measures of well-being. Initially a handful of studies attempted to associate specific candidate genes (e.g. *5-HTTLPR*, *MAOA*, *FAAH*) to hedonic well-being ^29–32^. However, these studies were most likely underpowered and results have not been replicated. More recent molecular genetic approaches revealed that 5-10% of the variation in responses to single-item survey hedonic measures (happiness) is accounted for by genetic variants measured on presently used genotyping platforms ^33^. Additionally, a recent large genome-wide association study (GWAS; *N* = 298,420) identified the first three genetic variants (two at chromosome 5 (rs3756290 and rs4958581) and one at chromosome 20 (rs2075677)) associated with SWB, defined as a combination of hedonic measurements like happiness and satisfaction with life ^34^.

There have only been two attempts to use molecular genetic data to reveal the overlap and distinction between hedonic and eudaimonic well-being ^35, 36^. The first study showed divergent transcriptome profiles between both measurements ^35^. Hedonic well-being was associated with up-regulated gene expression of a conserved transcriptional response to adversity (CTRA), while eudaimonic well-being was associated with CTRA down-regulation. After substantial critiques and replies ^37–40^, the authors of the initial finding replicated part of the results by showing a significant inverse relation between down-regulated CTRA expression and eudaimonic well-being ^36^. Based on these results, the authors conclude that eudaimonic well-being might play a more significant role in the link between well-being and health, than hedonic well-being.

The availability of large-scale molecular data make it possible to gain more insight into the genetic factors underpinning overlap and distinction between hedonic and eudaimonic well-being. In the current paper, we therefore leverage data from the UK Biobank and estimate the molecular genetic based heritability and bivariate genetic correlation. To this end, we conduct the first genome-wide association study (GWAS) to identify genetic variants associated with eudaimonic well-being as well as a GWAS for hedonic well-being. As the genetic architecture can be a reflection of common biology, we annotate the genome-wide association results using gene-mapping and tissue specific enrichment analyses. Additionally, the genetic correlation can be a product of a causal relationship between hedonic –and eudaimonic well-being. Therefore, using a bi-directional two-sample Mendelian Randomization (MR) design, we assess the direction of the relationship between hedonic and eudaimonic well-being. Finally, we estimate whether hedonic and eudaimonic Well-being show different genetic correlations patterns with positively and negatively related traits.

## Results

### Descriptive statistics and phenotypic correlation

For eudaimonic well-being, females and males mean scores were similar (mean =3.69, sd = 0.82 and 0.83, t = -0.79, *P* = 0.43). For hedonic well-being, males were significantly, but only slightly, happier (mean 4.52, sd = 0.74) than females (mean 4.51, sd =0.72) (t = 4.00, *P* < 0.001). Eudaimonic and hedonic well-being were moderately correlated (*r = 0.53, P* < 0.001; **Figure 2**).

**Figure 1:**
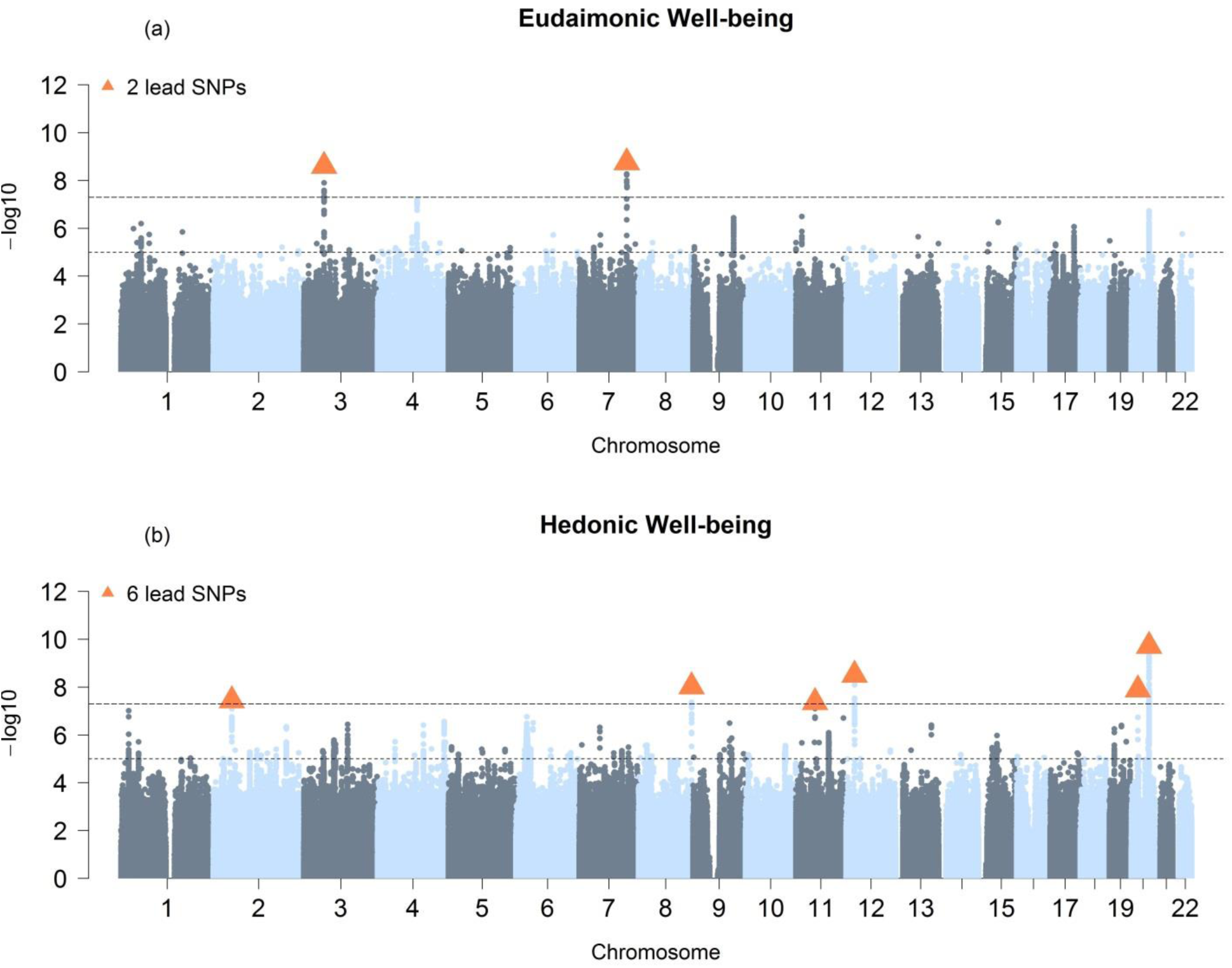
Manhattan Plot for GWAS results. Result is shown for **(a)** Univariate GWAS of eudaimonic well-being and, **(b)** N-weighed GWAMA of hedonic well-being. The *x* axis shows chromosomal position, and the *y* axis shows association significance on a −log10 scale. The upper dashed line marks the threshold for genome-wide significance (*P* = 5×10^−8^), and the lower dashed line marks the threshold for nominal significance (*P* = 1×10^−5^). Each approximately independent genome-wide significant association (lead SNP) is marked by an orange **Δ**. Each lead SNP is the SNP with the lowest *P* value within the locus, as defined by our clumping algorithm

**Figure 2:**
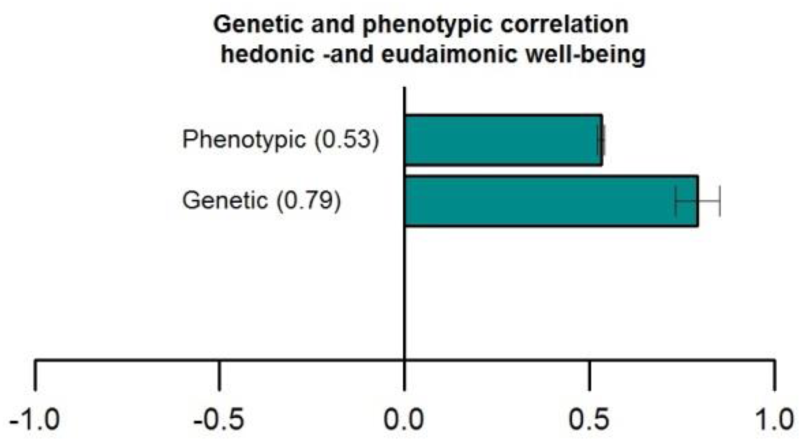
Phenotypic and genetic correlations between eudaimonic and hedonic well-being with their corresponding 95% confidence intervals.

### Genome-wide association analyses

For eudaimonic well-being, 2 genetic variants reached genome-wide significance (**Table 1 and Figure 1A**). The two univariate GWAS for hedonic well-being (UKB ID 4526 and UKB ID 20458) identified, respectively 1 and 2 genome-wide significant hits (**Supplementary Table 2 and Supplementary Figure 1-2**). The genomic inflation factor (lamda Genomic Control) of eudaimonic well-being (λ_GC_ = 1.14) and hedonic well-being (λ_GC__UKB ID 4526 = 1.13 and λ_GC__UKB ID 20458 = 1.13) were inflated. The estimated intercept from LD Score regression, though, did not exceed 1.02, indicating that nearly all the inflation is the GWAS analyses is due to polygenic signal rather than bias ^41^ (**Supplementary Table 3**). The multivariate N-weighted GWAMA for the two hedonic GWAS analyses yielded 6 genetic variants for hedonic well-being that reached genome-wide significance (λ_GC_ = 1.21, LD intercept = 1.00; **Figure 1B, Table 1, and Supplementary Table 3**). The significant SNPs associated with eudaimonic well-being had low *P*-values (7.6×10^-4^ for rs7618327 and 3.4×10^-5^ for rs7618327) in the hedonic analyses. Three out of 6 significant SNPs associated with hedonic well-being had low *P*-values (*P* < 3.6 x 10^-5^) in the eudaimonic GWAS.

**Table 1:** Genome-wide significant hits for eudaimonic -and hedonic well-being.

| **Eudaimonic well-being** |  |  |  |  |  |  |  |  |  |  |  |
| --- | --- | --- | --- | --- | --- | --- | --- | --- | --- | --- | --- |
| **SNP** | **RS** | **CHR** | **BP** | **A1** | **A2** | **Z** | **P** | **N** | **EAF** | **BETA** | **SE** |
| 7:127671511 | rs79520962 | 7 | 127671511 | A | G | -6.015 | 1.80E-09 | 108154 | 0.05 | -0.051 | 0.009 |
| 3:54376990 | rs7618327 | 3 | 54376990 | G | A | -5.961 | 2.52E-09 | 108154 | 0.12 | -0.033 | 0.006 |
| **Hedonic well-being** |  |  |  |  |  |  |  |  |  |  |  |
| **Multivariate** |  |  |  |  |  |  |  |  |  |  |  |
| 20:47746974 | rs34841991 | 20 | 47746974 | C | T | 6.367 | 1.92E-10 | 221575 | 0.24 | 0.022 | 0.004 |
| 12:22874365 | rs261909 | 12 | 22874365 | C | G | 5.925 | 3.12E-09 | 221575 | 0.44 | 0.018 | 0.003 |
| 8:142617261 | rs746839 | 8 | 142617261 | G | C | -5.739 | 9.53E-09 | 221575 | 0.38 | -0.018 | 0.003 |
| 20:17445078 | rs4239724 | 20 | 17445078 | G | A | -5.689 | 1.28E-08 | 221575 | 0.22 | -0.021 | 0.004 |
| 2:49222872 | rs6732220 | 2 | 49222872 | C | G | 5.506 | 3.68E-08 | 221575 | 0.77 | 0.020 | 0.004 |
| 11:51477511 | rs146213057 | 11 | 51477511 | A | G | 5.476 | 4.36E-08 | 221575 | 0.01 | 0.084 | 0.015 |

CHR = chromosome, BP= Base Pair, A1 = Effect allele, A2 = Other allele, Z = Zscore, P = P-value, N = sample size, EAF = Estimated Allele Frequency, SE = Standard Error

### SNP heritability and Genetic Correlation

For eudaimonic well-being, SNP h^2^ was 6.2% (se = 0.005), while for hedonic well-being the SNP h^2^ was 6.2% (se = 0.005) (UKB ID 4526) and 6.4% (se =0.005) (UKB ID 20458; **Supplementary Table 3**). The genetic correlation between the two measurements of hedonic wellbeing was –as expected-extremely high (0.99, *P* < 0.001). Additionally, the genetic correlation between eudaimonic and hedonic well-being was *r*_g_ = 0.78, (*P* < 0.001, **Figure 2 and Supplementary Table 4**).

### Polygenic prediction

Polygenic scores were calculated for 10 *P*-value thresholds, using Caucasian UK Biobank participants with non-British ancestry as an independent sample. PRS based on the hedonic well-being GWAMA explained 0.83% (*P* = 2.81×10^-18^) of the variance in eudaimonic well-being whereas PRS based on the eudaimonic well-being GWAS explained 0.43% (*P* = 2.60×10^-10^) of the variance in hedonic well-being. A complete overview of the polygenic scores including all thresholds can be found in **Supplementary Table 5 and Supplementary Figure 3.**

### Functional annotation

#### Eudaimonic well-being

We searched the NHGI GWAS catalog to determine which of the lead SNP (*P* < 5×10^-8^, independent from each other at *r^2^*< 0.1) associated with eudaimonic well-being have been previously reported. This search initially revealed that none of the variants are previously reported. However, if we look at the results of the gene-based test as computed by MAGMA including all SNPs with a P value below 0.05, genes associated with Educational attainment ^42^ (*ARFGEF2*), Subjective Well-being ^34^ (*ARFGEF2, CSE1L*) and height ^43^ (*STAU1, ZFAS1*) were found.

Based on the eudaimonic well-being GWAS, 3 genes were found through positional mapping, 1 through eQTL mapping, and 13 through chromatine interaction-mapping (**Supplementary Tables 6-8**). Looking at the results of the gene-based test as computed by MAGMA including all SNPs with a *P* value below 0.05, 10 genes were associated with eudaimonic well-being (**Supplementary Table 9**). Of these 27 genes in total, one gene (*SND1*) was implicated in all four methods. The *SND1* gene encodes a transcriptional co-activator that interacts with the acidic domain Epstein-Barr virus nuclear antigen (EBNA 2), a transcriptional activator that is required for B-lymphocyte transformation. Proteins encode by this gene are thought to be essential for normal cell growth (https://www.ncbi.nlm.nih.gov/gene/27044).

#### Hedonic well-being

We first searched the NHGI GWAS catalog to determine which of the lead SNP associated with hedonic well-being have been previously reported. Here we found that the variants have been reported in Educational attainment ^42^ (*ARFGEF2*), Obesity-related traits ^44^ (*PCSK2, ARFGEF2*), Subjective Well-being ^34^ (*ARFGEF2, CSE1L*) and height ^43^ (*STAU1, ZFAS1*) (**Supplementary Table 10**).

Based on the multivariate N-weighted GWAMA, 7 genes were implicated through positional mapping, 9 through eQTL mapping, and 50 through chromatine interaction-mapping (**Supplementary Tables 11-13**). Using the results of the gene-based test as computed by MAGMA including all SNPs with a P value below 0.05, 35 genes were associated with hedonic well-being (**Supplementary Table 14**). Of these 101 genes in total, 16 were found in more than one strategy. Of these, two genes (*CSE1L*, *STAU1*) were implicated by all four methods. Proteins encode by *CSE1L*, may play a role in apoptosis and in cell proliferation (https://www.ncbi.nlm.nih.gov/gene/1434?otool=inlvulib). The *STAU1* gene is a member of the family of double stranded RNA (dsRNA)-binding proteins involved in the transport and/or localization of mRNAs to different subcellular compartments. *STAU1* contains a microtubule-binding domain similar to that of microtubule-associated protein 1B (*MAP1B*) and bind tubulin (https://www.ncbi.nlm.nih.gov/gene/6780).

#### Tissue Specific expression

Tissue expression analysis, performed on GTEx RNA-sq data, showed significant enrichment in the brain cortex, brain cerebellum, frontal cortex, as well as the cerebellar hemisphere for eudaimonic well-being. In contrast, no significant results were found for hedonic well-being, although brain tissues were top ranked in their enrichment (**Supplementary Table 15, 16, Supplementary Figure 4**).

#### Mendelian Randomization

To test the direction of the relationship between hedonic and eudaimonic well-being a Two-Sample Mendelian Randomization (2S-MR) design was applied. We found little to no evidence for a causal effect of eudaimonic well-being on hedonic well-being using either suggestive SNPs (*P* < 1 × 10^-5^) or genome-wide significant SNPs (*P* < 5 × 10^-8^) as genetic instruments. Only the weighted median regression was significant (*P* = 0.0117; **Supplementary Tables 17A, 18**). More support was found for a causal effect of hedonic well-being on eudaimonic well-being. Using suggestive SNPs (*N = 78)* all included methods were Bonferroni significant (*P* < 0.0125) except the MR-Egger results (**Supplementary Table 17B,C**). However, sensitivity tests indicated the presence of some horizontal pleiotropy (**Supplementary Table 19**). Using genome-wide significant SNPs (*N* = 6) as genetic instruments, the simple mode, weighted median –and IVW regression were significant with the latter two surviving Bonferroni correction. The weighted mode and MR-Egger results were not significant). It is, however, known that the power to detect causal relationships using the MR-Egger estimator with few genetic instruments –as in our case- is low ^45^. In addition, sensitivity tests showed little to no bias to horizontal pleiotropy or heterogeneity (**Supplementary Table 19**). The Steiger test showed that the genetic instruments explained more variation in the exposure compared to the outcome suggesting a correct causal direction (**Supplementary Table 19**).

#### Genetic Correlations

Another way to study the relationship between eudaimonic and hedonic well-being is by comparing their genetic correlation patterns with positive and negative related traits. Overall we found a similar pattern for both eudaimoninc and hedonic well-being. Both were positively correlated with satisfaction with health (*rgEUD* = 0.53, *rgHED* = 0.61), financial satisfaction (*rgEUD* = 0.39, *rgHED* = 0.49), friendship satisfaction (*rgEUD* = 0.68, *rgHED* = 0.81), family Satisfaction (*rgEUD* = 0.65, *rgHED* = 0.76) and job satisfaction (*rgEUD* = 0.73, *rgHED* = 0.84). Negative correlations were found for irritable (*rgEUD* = -0.25, *rgHED* = -0.36), loneliness (*rgEUD* = -0.45, *rgHED* = -0.56), depressive symptoms (*rgEUD* = -0.32, *rgHED* = - 0.53), depression diagnosed by doctor (*rgEUD* = -0.37, *rgHED* = -0.51), and neuroticism (*rgEUD* = -0.45, *rgHED* = -0.58; **Figure 3 and Supplementary Table 20**). These similar patterns support the finding of a large genetic overlap between eudaimonic and hedonic well-being.

**Figure 3:**
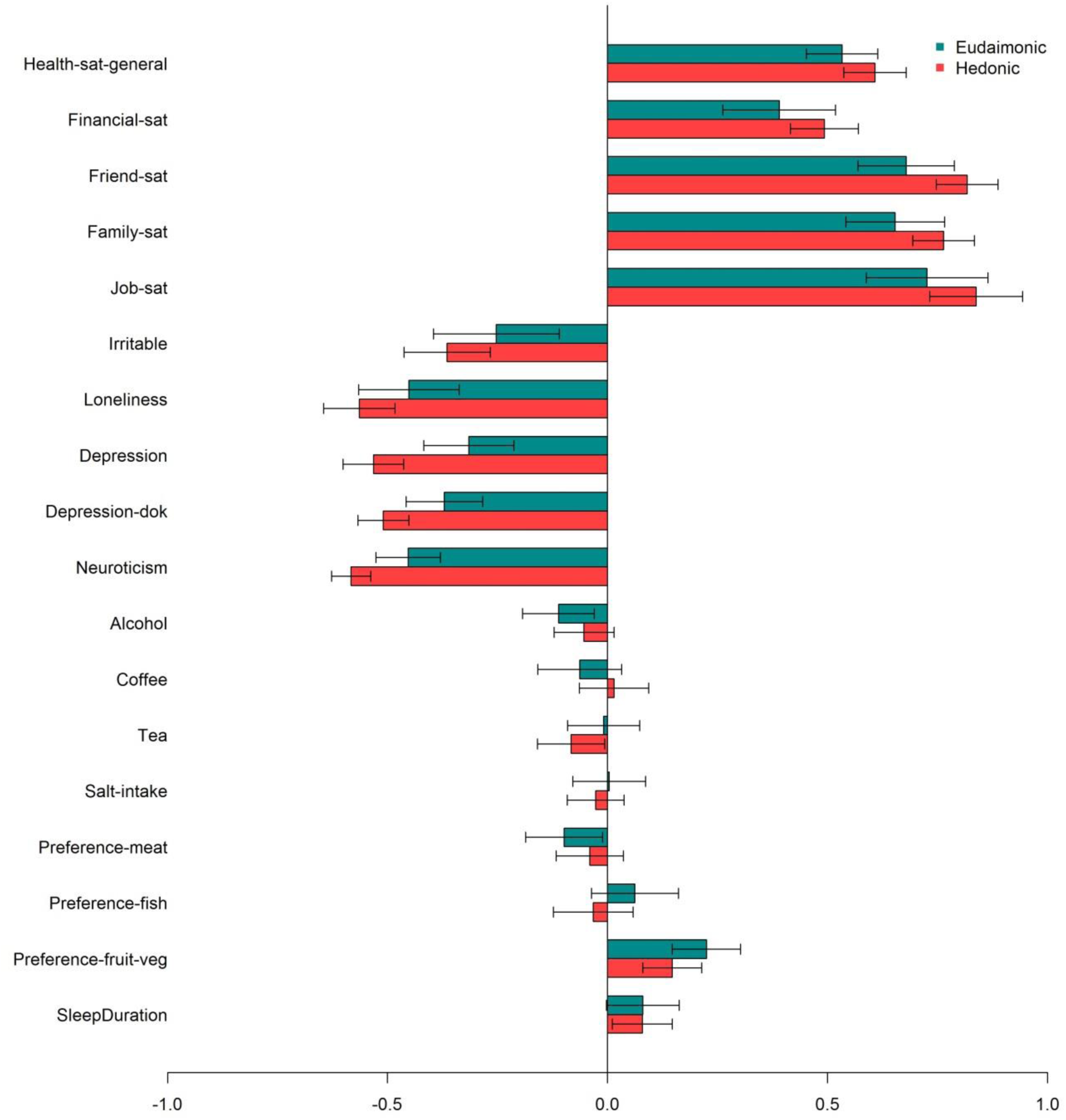
Genetic correlations between eudaimonic (blue) –and hedonic well-being (red) with (from top to bottom): satisfaction with health, financial satisfaction, friendship satisfaction, familial satisfaction, job satisfaction, irritable, loneliness, depression, depression diagnosed by a doctor, neuroticism, alcohol use, coffee use, tea use, salt intake, meat preference, fish preference, fruit preference and sleep duration. 95% confidence intervals are provided.

## Discussions

In this article, we provide evidence for a strong genetic overlap between hedonic and eudaimonic well-being. Our analyses revealed a moderate phenotypic correlation (*r* = 0.53), but a high genetic correlation (*r_g_* = 0.78). Our results include the first two genome-wide significant independent loci for eudaimonic well-being and 6 independent loci for hedonic well-being. Biological annotation points to a central role for the central nervous system in both forms of well-being. Loci regulating expression showed significant enrichment in the brain cortex, brain cerebellum, frontal cortex, as well as the cerebellar hemisphere for eudaimonic well-being. No significant enrichment for hedonic well-being is observed, although brain tissues were top ranked.

The high genetic correlation between the two forms of well-being can be a product of a causal relationship between the two traits. We report evidence for a causal effect of hedonic well-being on eudaimonic well-being whereas no evidence was found for the reverse. Although the findings are strengthened by the two-sample MR approach, which has considerable higher power to pick up causal effects compared to the one-sample MR approach, these results should be interpreted with caution. The lack of evidence for a causal effect of eudaimonic well-being on hedonic well-being might be explained by limited power of the chosen instrumental variable, the polygenicity of both measures, or it may be the results of violation of the MR design ^46^. In addition, sensitivity analyses revealed the presence of horizontal pleiotropy using suggestive SNPs as genetic instruments, indicating that these results might be driven by an unknown third variable.

Further evidence for a shared genetic architecture between hedonic and eudamonic well-being is provided by the similar patterns of genetic correlations with other traits. Largest correlations were found for job satisfaction followed by friendship –and family satisfaction and general health satisfaction. Remarkably, in contrast to job satisfaction, financial satisfaction showed the lowest correlation with both eudaimonic – and hedonic well-being. Although, hedonic well-being showed larger point estimates with all positive related phenotypes, the 95% confident intervals were overlapping. Genetic correlations with negative related phenotypes were for both measures largest for neuroticism followed by loneliness, depression (2X) and irritable. Similar to positive related traits, genetic correlations patterns were similar, except for neuroticism and depression, that were more strongly correlated with hedonic well-being.

Besides adding to the ongoing debate on the overlap and distinction between hedonic and eudaimonic well-being the current study provides novel insight into the genetics of well-being by identifying genome-wide significant genetic variants that explain differences in eudaimonic well-being. These variants have not been associated with a complex trait before, and thus warrant replication. Robustness of the current findings, though, is reflected in our hits for hedonic well-being. These genome-wide significant hits have been found to be associated with hedonic well-being in an earlier large scale study ^34^.

The findings of this study should be interpreted in light of the following limitations. One is that eudaimonic and hedonic well-being are based on single item measurements. We, however, have explicitly chosen not to include all other available hedonic results of our previous work ^34, 47^, to leverage the power of homogeneity of the UK Biobank dataset and to ease the interpretation of our findings. Research studying higher-quality measures of the various facets of well-being is a critical next step. Our results can help facilitate such work because, if the variants we identify are used as candidates, studies conducted in the smaller samples in which more fine-grained phenotype measures are available can be well powered.

In conclusion, while we found a moderate phenotypic correlation between eudaimonic and hedonic well-being, we report a strong genetic correlation. Future studies should acknowledge the strong genetic correlation between eudaimonic and hedonic well-being and include both to increase our understanding of the (genetic) etiology of well-being.

## Methods

### Participants

We analyzed data from the UK Biobank project ^48^. The UK Biobank is a prospective study designed to be a resource for research into the causes of disease in middle and old age. The study protocol and information about data access are available online (http://www.ukbiobank.ac.uk/wp-content/uploads/2011/11/UK-Biobank-Protocol.pdf) and more details on the recruitment and study design have been published elsewhere ^48^. The UK Biobank study was approved by the North West Multi-Centre Research Ethics Committee (reference number 06/ MRE08/65), and at recruitment all participants gave informed consent to participate in UK Biobank and be followed-up, using a signature capture device. All experiments were performed in accordance with guidelines and regulations from these committees. In brief, all participants were registered with the UK National Health Service (NHS) and lived within 25 miles (40 km) of one of the assessment centres. The UK Biobank invited 9.2 million people to participate through postal invitation with a telephone follow-up, with a response rate of 5.7%. A total of 503,317 men and women aged 40–70 years were recruited in assessment centres across England, Wales and Scotland, between 2006 and 2010. In total, 608 participants have subsequently withdrawn from the study and their data were not available for analysis. Participants attended 1 of 22 assessment centers across the UK, at which they completed a touch-key questionnaire, had a face-to-face interview with a trained nurse, and underwent physical assessments. Participants completed sociodemographic questionnaires, which included questions on financial satisfaction and income as well as questionnaires about their physical and mental health.

Data access permission was granted under UKB application 25472 (PI Bartels). For the discovery genome-wide association analyses we used data of ≍ 110K UK-habitant Caucasian individuals only. A full overview of the included participants with valid phenotypic measurements as well as genetic data is presented in

## Supplemental Table 1

### Phenotypic data

Eudaimonic well-being was assessed in the online follow-up with its core element meaning in life (“To what extent do you feel your life to be meaningful?”; UKB Data-Field 20460). Answers were provided on a 5-item likert scale that ranged from “Not at all” (score 1) to “An extreme amount” (score 6). Information on eudaimonic well-being and genotypic data were available for 108,154 UK Biobank participants (56% female).

Hedonic Well-being was assessed with its core element general happiness (“In general how happy are you?”; UKB Data-Field 4526 & UKB Data-Field 20458). Answers were provided on a 6-item likert scale that ranged from “Extremely happy” (score 1) to “Extremely unhappy” (score 6). Scores were reversed so that a higher score was associated with higher levels of happiness. Hedonic well-being, as part of the touchscreen questionnaire on psychological factors and mental health (data-field 4526), was available for 111,470 individuals. Hedonic well-being was also assessed in the online follow-up (data-field 20458) and this measure is available for 110,105 individuals. Almost forty thousand individuals (n=39,999) participated in both assessments. In total, information on hedonic well-being and genotypic data were was available for 181,578 UK Biobank participants (49% female).

### Genotypic data

Participants were genotyped using one of two platforms: The affymetrix UK BiLEVE Axiom array or the Affymetrix UK Biobank Axiom array. The genetic data underwent rigorous quality control and was phased and imputed against a reference panel of Haplotype Reference Consortium (HRC), UK10K and 1000 Genomes Phase 3 haplotypes^49^. Due to an issue with the imputation of UK10K and 1000 Genomes variants, analyses were restricted to HRC variants only. Samples were excluded based on the following genotype-based criteria; non-European ancestry, relatedness, mismatch between genetic sex and self-reported gender, outlying heterozygosity, and excessive missingness ^49^. For more details on the UK Biobank genotyping, imputation, and quality control procedures see ^50^.

### Descriptive statistics and phenotypic correlation

Descriptive statistics and spearman’s rank correlation between eudaimonic and hedonic well-being were calculated in R. We, furthermore, tested for sex and age effects on mean levels.

### Univariate Genome-wide association analyses

Univariate genome-wide association analyses for eudaimonic well-being and for hedonic well-being (touchscreen measure and online follow-up separately) were performed in PLINK ^51, 52^ using a linear regression model of additive allelic effects. Standard pre-GWAS-quality control filters were applied, which included removing SNPs with minor allele frequency < 0.005 and/or with an INFO-score < 0.8 for imputed SNPs, and removing individuals with ambiguous sex and/or non-British ancestry. We, furthermore, randomly selected 1 individual from each closely related pair (i.e. parent offspring pairs, sibling pairs). The GWAS included 40 principal components, age, sex, and a chip dummy as covariates. Additionally, following a pre-specified analysis plan, we conducted a stringent post-GWA quality control (QC) protocol based on the paper of Winkler and colleagues ^53^.

### Multivariate Genome-wide association analyses

To increase the effective sample size, we conducted multivariate N-Weighted genome-wide association meta-analyses (GWAMA) by leveraging the association between the two hedonic well-being univariate GWAS analyses (UKB Data-field 4526 and 20458, n_obs_ total = 221,575). The dependence between effect sizes (error correlation) induced by sample overlap in both these GWAMAs was estimated from the genome-wide summary statistics of the univariate GWAS analyses using LD score regression ^54^. Knowledge of the error correlation between the univariate GWAS analyses allowed us to meta-analyze them together, providing a gain in power while guarding against inflated type I error rates. For a detailed description on performing N-weighted GWAMA, please see Baselmans and colleagues ^47^.

### SNP heritability and Genetic Correlation

SNP heritability for eudaimonic and hedonic well-being separately was estimated using bivariate LD Score Regression ^54, 55^. The same methodology was used to estimate the genetic correlation between the two measures of hedonic well-being and between eudaimonic and hedonic well-being. LD scores regression produces unbiased estimates even in the presence of sample overlap and only requires summary statistics and a reference panel from which to estimate each SNP’s “LD score” (the amount of genetic variation tagged by a SNP). We used the file of LD scores computed by Finucane et al. ^56^ using genotypic data from a European-ancestry population (see https://github.com/bulik/ldsc/wiki/Genetic-Correlation, accessed September 8, 2017).

### Polygenic prediction

We performed polygenic risk score prediction (PRS) using Caucasian UK Biobank participants with non-British ancestry as independent prediction sample (n_obs_ = 28,582). For eudaimonic well-being, polygenic prediction was performed in 9,088 individuals. For hedonic well-being, we used phenotypic measurements closest to genotype-collection (UKB Data-Field 20458) for polygenic scores and scores were available for 9,276 individuals. The weights used for the polygenic scores are based on the univariate GWAS (eudaimonic) and multivariate GWAMA (hedonic well-being). Polygenic scores were based the genotyped SNPs (n_obs_ = 619,049). To calculate the incremental R^2^, the phenotypes (eudaimonic and hedonic well-being) were standardized and regressed on sex and age as well as principal components, which were included to correct for ancestry. Next, the same analysis was repeated with inclusion of the polygenic scores. The differences in R^2^ between both regression is referred to as incremental R^2^. To obtain 95% confidence intervals (CI) around the incremental R^2^ ’s, bootstrapping was performed with 2000 repetitions.

### Functional annotation

Functional annotation was performed in FUMA ^57^ (http://fumactglab.nl) for the eudaimonic well-being GWAS and the hedonic well-being GWAMAs. Lead SNPs were defined as having a genome-wide significant P values (5×10^-8^) and being independent from each other (r^2^< 0.1). Functional annotation was performed on these lead SNPs and SNPs with P < 0.05, MAF < 0.01, and in high LD (r^2^ > 0.6) with those lead SNPs.

### Gene-mapping

This set of SNPs was mapped to genes in FUMA using three strategies. The SNPs were mapped to genes based on 1) their physical distance (i.e. within 10kb window), 2) significant eQTL association (i.e. the expression of that gene is associated with allelic variation at the SNP). eQTL mapping in FUMA uses information from the GTEx, Blood eQTL browser, and BIOS QTL browser, and is based on cis-eQTLs that can map SNPs to genes up to 1MB apart. A false discovery rate (FDR) of 0.05 was applied to define significant eQTL associations. 3) a significant chromatin interaction between a genomic region and promoter regions of genes (250bp up and 500bp downstream of transcription start site (TSS)). Chromatine interaction mapping can involve long-range interaction as it does not have a distance boundary as in eQTL mapping. We used a FDR p-value of 1×10^-5^ to define significant interactions.

Finally, given our modest sample size and expected polygenicity of our phenotypes, we added an extra strategy in which all SNPs (P < 0.05) were included and mapped to genes based on physical distance (i.e. within 10kb window) from known protein coding genes (GRCh37/hg19). Genome-wide significance for this test was defined at P = 0.05/ 18187 = 2.74×10^-6^.

### Tissue Expression Analysis (MAGMA)

To test the relationship between highly expressed genes in a specific tissue and genetic associations, gene-property analysis is performed using average expression of genes per tissue type as a gene covariate. Gene expression values are log^2^ transformed average RPKM (Reads Per Kilobase Million) per tissue type after winsorized at 50 based on GTEx RNA-seq data. Tissue expression analysis is performed for 53 specific tissue types separately. The result of the gene analysis (gene-based P value) were used in MAGMA to test for one-side increased expression conditioned on average expression across all tissue types.

### Mendelian Randomization

To gain insight whether eudaimonic well-being causes hedonic well-being or the other way around we applied bi-directional two-sample MR (2S-MR) ^58–60^. This method takes the genetic instrument from a GWA study on the exposure variable (gene-exposure association) and then identifies the same genetic instrument in a GWA study on the outcome variable (gene-outcome association). MR is based on the following three assumptions; 1) the genetic instrument is predictive of the exposure variable, 2) the genetic instrument is independent of confounders, and 3) there is no pleiotropy (the genetic instrument does not affect the outcome variable, other than that through its possible causal effect on the exposure variable). If these assumptions are met, genetic variants that predict the exposure variable should through the causal chain, also predict the outcome variable.

All analyses were conducted with the 2SMR package of MR-Base ^61^ in R (https://github.com/MRCIEU/MRInstruments). Genetic instruments were constructed using two p-value thresholds; 1) where only genetic variants that exceeded the threshold for genome-wide significance (P < 5×10^-8^) were included and 2) where genetic variants that exceeded a less stringent threshold (P < 1×10^-5^) were included. A priori to MR analyses, genetic variants were pruned r^2^< 0.001) and if needed proxies were identified (r^2^ ≥ 0.8).

When a genetic instrument consists of less than three genetic variants, the causal effect was estimated using the Wald ratio method, which is computed as the outcome association divided by the exposure ^62^. Additional, a maximum-likelihood test was performed for the combined test.

When the genetic instrument consisted of more than 2 genetic variants, we used the following five methods: 1) MR-Egger regression, which relaxes the assumption that the effects of the variants on the outcome are entirely mediated via the exposure. MR-Egger allows for each variant to exhibit some pleiotropy, but assumes that each gene’s association with the exposure is independent in magnitude from its pleiotropic effects (the InSIDE assumption ^45^. 2) We conducted weighted median regression analyses, which provided a consistent estimate for the true causal effect when up to half of the weight in the MR analysis pertains from genetic variants that exert pleiotropic effects on the outcome^63^. 3) The Inverse-Variance Weighted (IVW) linear regression was applied, which sums the ratio estimates of all variants in a weighted average formula ^60^. 4) The Simple mode and 5) the Weighted mode that offers robustness to horizontal pleiotropy relying on the fundamental assumption termed the ZEro Modal Pleiotropy Assumption (ZEMPA). This assumption states that in large sample sizes, the largest subset of variants with the same ratio estimate comprises the valid instruments. Invalid instruments may have different ratio estimates asymptotically, but there is no larger subset of invalid instruments with the same ratio estimate than the subset of valid instruments ^64^

Some of the MR tests can also perform sensitivity tests to investigate the robustness of the obtained results. Tests for heterogeneity were performed using MR-Egger and the Invers-Variance Weighted regression. To identify horizontal pleiotropy, the value of the MR-Egger intercept provides an indication of the degree of pleiotropy affecting the results. Finally, after each analysis, a Steiger test ^65, 66^ was performed to make sure that the chosen instrument influence the exposure first and then the outcome through the exposure. To this end, the Steiger test calculated the variance explained (R^2^) in the exposure and in the outcome by the instrumenting SNP and subsequently test whether the variance in the outcome is less than in the exposure.

As our eudaimonic –and hedonic well-being measurements were both part of the UK Biobank project, sample overlap was inevitable. However, when sample overlap is present in two-sample MR, results can be biased with increased Type 1 error rates, that are linearly related to the proportion of sample overlap ^67^ To overcome this bias, we used the following strategy: Hedonic well-being data collection in the UK Biobank was done at two different time points. For both measurements, we calculated the sample overlap with eudaimonic hedonic well-being. We found that hedonic well-being (UK Biobank ID 4526) had 72,146 unique participants compared to the eudaimonic well-being. Therefore, we performed a separate univariate GWAS for this group and performed MR analyses using the two strategies described above. For genetic instruments (significant and suggestive SNPs), we used the results of the multivariate GWAMA of hedonic well-being and extracted these from the hedonic well-being GWAS including only the unique participants relative to the Eudaimonic GWAS.

Since we tested four different causal associations, we considered an alpha level of 0.0125 (0,05/4) as Bonferroni significant.

### Genetic Correlation

To test whether hedonic or eudaimonic well-being are genetically different correlated with a set of related phenotypes, bivariate LD Score regression was applied with both measures of well-being and the following UK Biobank summary statistics: satisfaction with health (UKB ID 20459), financial satisfaction (UKB ID 4581), friendship satisfaction (UKB ID 4570), family satisfaction (UKB ID 4559), job satisfcation (UKB ID 4537), irritable (UKB ID 4653), loneliness (UKB ID 2020), depressive symtoms (UKB ID 2100), depression diagnosed by doctor (UKB ID 2090), neuroticism (UKB ID 20127), alcohol (UKB ID 1558), coffee (UKB ID 1498), tea (UKB ID 1488), salt (UKB ID 1478), food preference meat (UKB ID 1349), food preference fish (UKB ID 1329), food preference fruit/vegetarian (UKB ID 1289), sleep duration (UKB ID 1160). For every genetic correlation 95% confident intervals were calculated.

### Data Availability Statement

The datasets generated during and/or analysed during the current study are available from the corresponding author on reasonable request.

## Acknowledgments

M. Bartels and B.M.L Baselmans are financially supported by the University Research Chair position of M. Bartels. This research has been conducted using UK Biobank resource (application number 25472 (PI Bartels). We would like to thank the participants and researches who collected and contributed to the data. We would like to thank Michel Nivard, Matthijs van der Zee, and Hill Fung Ip for their contributions in processing the UK Biobank genetic data.

## Author Contribution

**M.B.** supervised the entire project. **M.B.** and **B.B.** wrote the main manuscript text. **M.B.** was responsible for data curation. **M.B.** and **B.B.** performed formal analyses. **B.B.** prepared all (supplementary) figures and tables. **M.B.** and **B.B.** took care of the project administration.

## Supplementary Figures

**Figure 1:** Manhattan plot for the univariate GWAS results of Hedonic Well-being (UKB ID 4526)

**Figure 2:** Manhattan plot for the univariate GWAS results of Hedonic Well-being (UKB ID 20458)

**Figure 3:** Polygenic scores including 10 *P*-value thresholds for Eudaimonic and Hedonic Well-being

**Figure 4:** Cell type specific enrichment for eudaimonic –and hedonic well-being

**Supplementary Figure 1:**
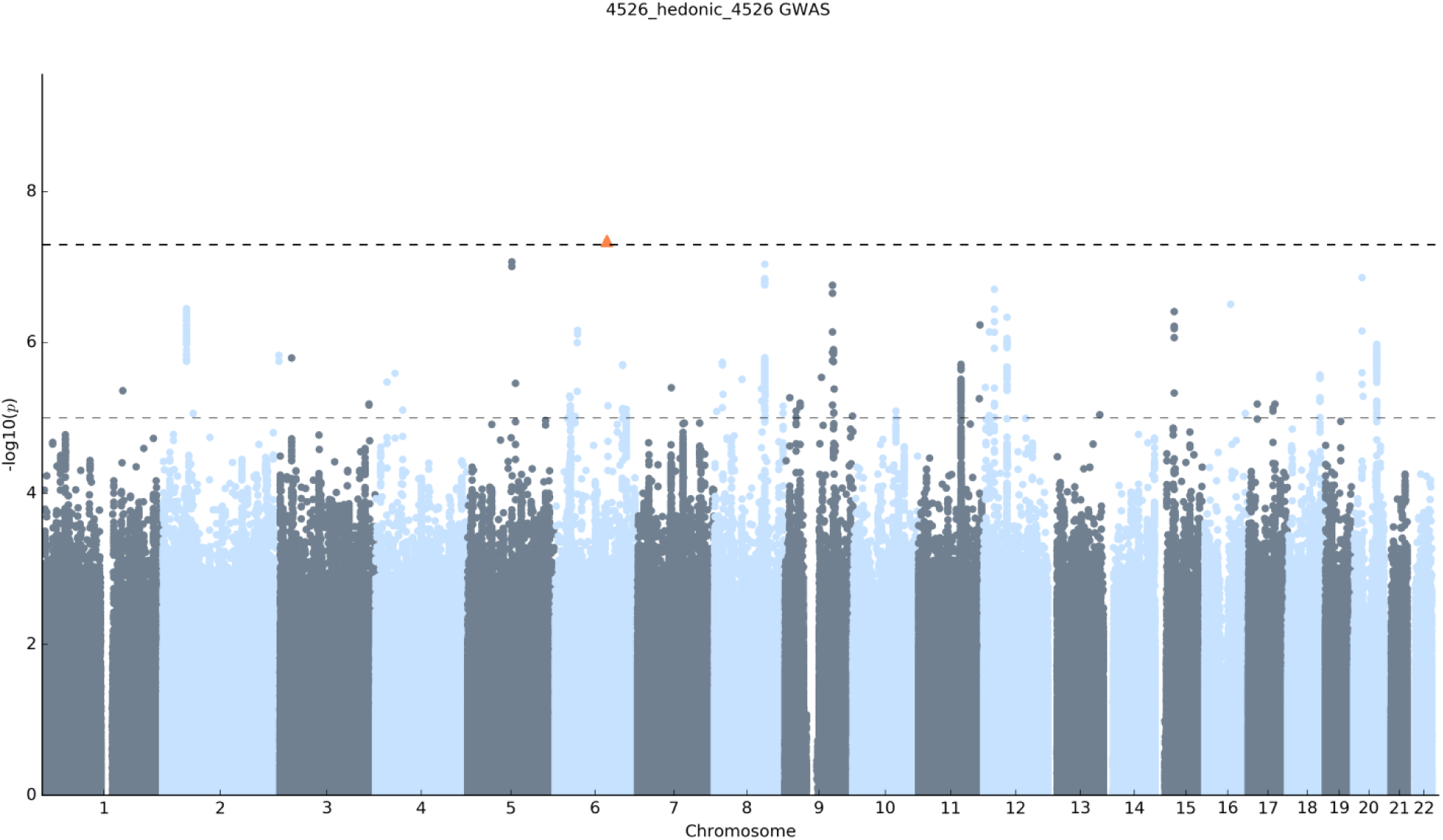
Manhattan Plot for GWAS results. Result is shown for Hedonic well-being (UKB ID 4526). The *x* axis shows chromosomal position, and the *y* axis shows association significance on a −log10 scale. The upper dashed line marks the threshold for genome-wide significance (*P* = 5 × 10−8), and the lower dashed line marks the threshold for nominal significance (*P* = 1 × 10−5). Each approximately independent genome-wide significant association (lead SNP) is marked by an orange **Δ**. Each lead SNP is the SNP with the lowest *P* value within the locus, as defined by our clumping algorithm

**Supplementary Figure 2:**
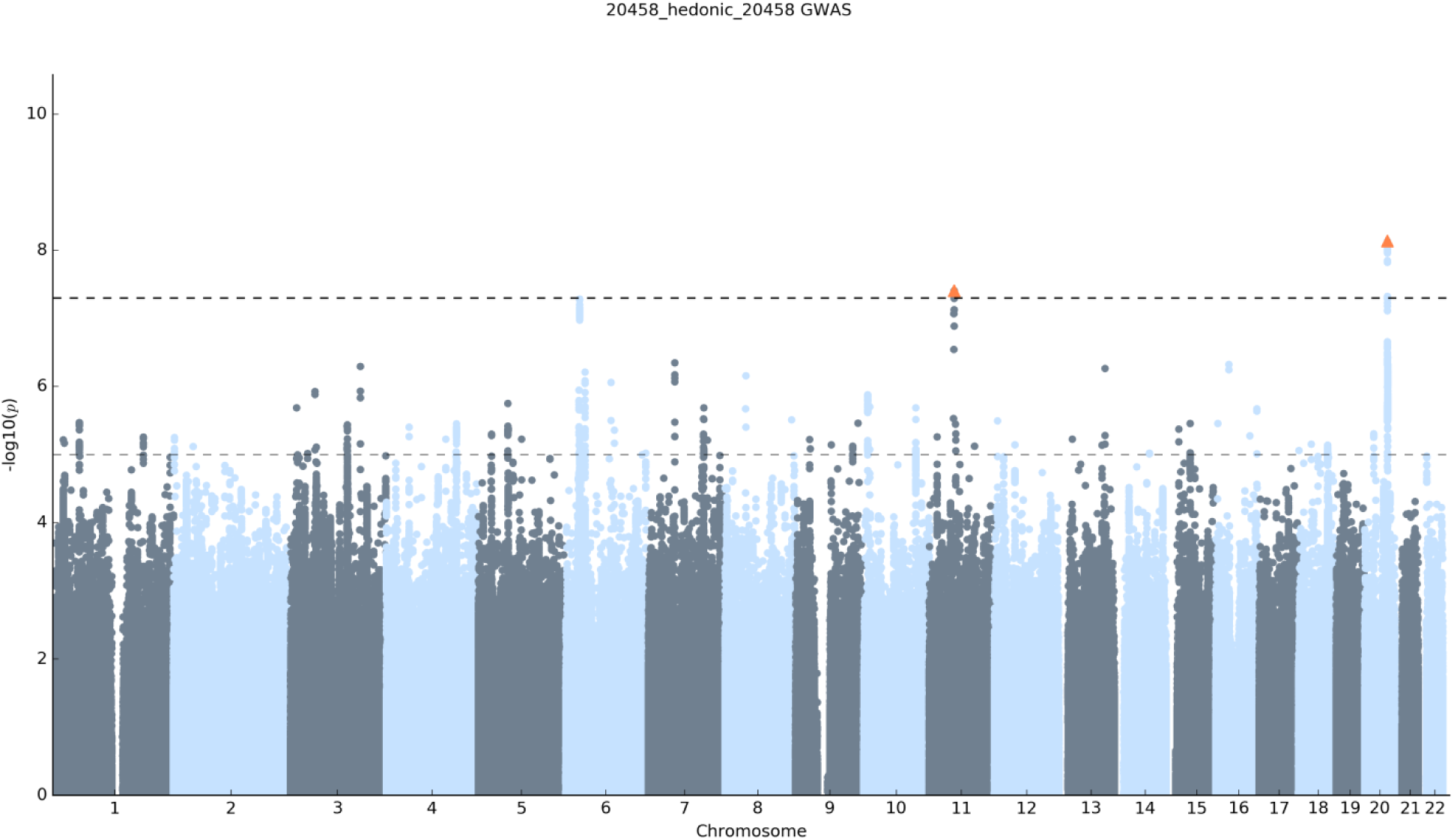
Manhattan Plot for GWAS results. Result is shown for Hedonic well-being (UKB ID 20458). The *x* axis shows chromosomal position, and the *y* axis shows association significance on a −log10 scale. The upper dashed line marks the threshold for genome-wide significance (*P* = 5 × 10−8), and the lower dashed line marks the threshold for nominal significance (*P* = 1 × 10−5). Each approximately independent genome-wide significant association (lead SNP) is marked by an orange **Δ**. Each lead SNP is the SNP with the lowest *P* value within the locus, as defined by our clumping algorithm

**Figure 3:**
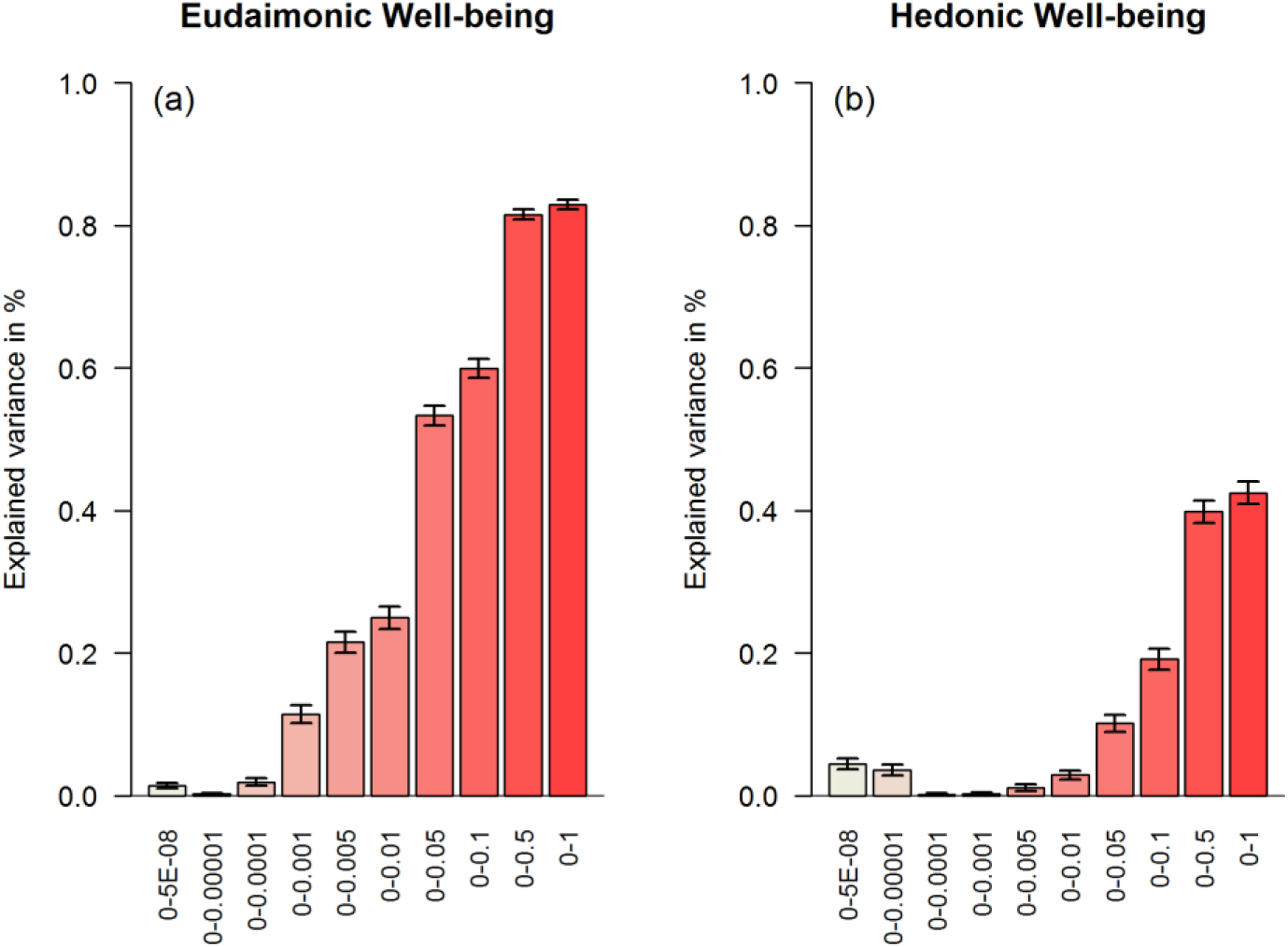
Polygenic scores using 10 different p-value thresholds for **(a)** Eudaimonic, and **(b)** Hedonic. The x axis shows the 10 different P value thresholds whereas the y axis shows the explained variance in percentage.

**Figure 4:**
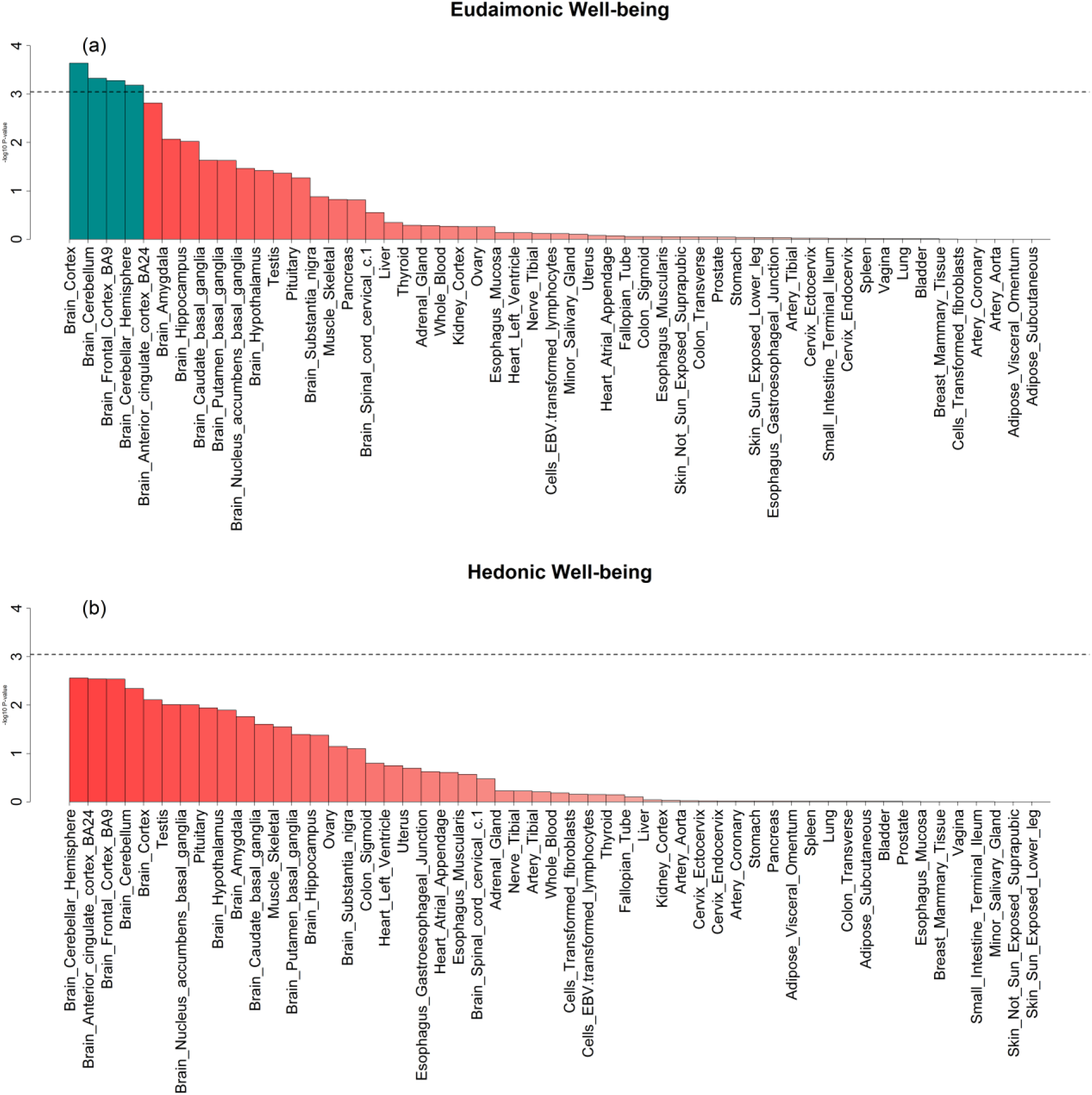
Tissue specific enrichment using 53 specific tissue types for **(a)** Eudaimonic, **(b)** Hedonic. Bar-graphs above the dashed line are significantly enriched. The x axis shows the 53 different categories whereas the y axis shows the –log^10^ P value. Bars in blue are significant enriched.

